# Experimental validation of computationally prioritized bisphosphonates reveals no direct *in vitro* antiviral activity against SARS-CoV-2

**DOI:** 10.64898/2026.09.07.749793

**Authors:** Mohammed Muzaffar-Ur-Rehman, Stephanie Olliff, Alexander J McAuley, Ashish Awasthi, Kondapalli Venkata Gowri Chandrasekhar, Sankaranarayanan Murugesan, Seshadri S Vasan

## Abstract

It is essential to experimentally evaluate antiviral efficacy predicted by molecular docking and related *in silico* approaches, which this study achieves for three bisphosphonates (alendronate, minodronate, zoledronate) and treprostinil, identified as potential antiviral therapies in our previous work. These investigational compounds had no detectable cytotoxicity at concentrations up to 25µM in Vero E6 cells but also failed to show any antiviral effect against the PQ.8.1 isolate of SARS-CoV-2, when compared to control drugs (ensitrelvir, nirmatrelvir and remdesivir). It appears that for bisphosphonates, any protective effect is likely due to other virus- or host-mediated mechanisms of action. This study demonstrates the importance of iterating between theoretical hypotheses and experiments to elucidate underlying mechanisms at play.

**Significance statement:** Repurposed drugs could have unexpected antiviral activities of their own as we previously demonstrated with fluvoxamine, therefore, experimentally ruling this out for bisphosphonates is important. Likely mechanisms of action could involve cathepsin and RdRp, then this could have wide implications beyond SARS-CoV-2. If bisphosphonates provide partial protection against COVID-19, then current and future vaccine and drug studies should take this into account in their design.

## Introduction

RNA viruses are responsible for many significant diseases, from Ebola, HIV and polio to respiratory diseases such as influenza, severe acute respiratory syndrome (SARS) and coronavirus disease 2019 (COVID-19) [1]. SARS coronavirus-2 (SARS-CoV-2), the causative agent of COVID-19, emerged in 2019 and rapidly evolved into a global pandemic resulting in over 779 million infections [2]. Since then, it has claimed approximately 7 million lives [3], resulted in an additional 2-fold excess deaths [4], caused severe socioeconomic disruption worldwide [2]. Although vaccination has substantially reduced severe outcomes, their effectiveness remains limited as the virus undergoes continuous evolution to adapt to antiviral immune responses [5]. Populations worldwide are experiencing vaccine distrust, fatigue and hesitancy [6–8], while high-risk groups such as immunocompromised individuals continue to remain vulnerable [9]. This means that COVID-19 and a future “Disease X” outbreak likely cannot be managed through vaccination alone. Accordingly, we need safe, effective and cost-effective treatment options in the medical countermeasure arsenal that can be used to treat these infections. Despite breakthroughs such as ensitrelvir and Paxlovid, the development of antiviral therapies is a long and complicated process. Moreover, it is important to understand whether any existing drug could enhance or interfere with COVID-19 medications.

Several direct-acting anti-viral drugs, such as remdesivir [10,11], molnupiravir [12], favipiravir [13], nirmatrelvir in combination with ritonavir (Paxlovid) [14] and ensitrelvir [15], have shown meaningful therapeutic benefit against COVID-19. However, these drugs are most effective when administered in the early stages of infection, leaving a critical void in the therapeutic timeline. These drugs also have limitations restricting their universal effectiveness in immunocompromised individuals, including transplant recipients, cancer patients, and individuals with immunosuppressive therapies, who are at increased risk for severe disease progression and poor vaccine responsiveness. For example, they can alter the plasma concentrations of immunosuppressive agents such as tacrolimus, everolimus, and cyclosporin; increasing the risk of toxicity or graft rejection [16–19]. Remdesivir, a nucleoside analogue and inhibitor of the viral RNA-dependent RNA polymerase (RdRp), requires intravenous administration, which limits its use in resource-constrained settings. It can exert significant strain on the kidneys especially in renal transplant recipients, causing renal toxicity [20,21]. It can lead to cardiovascular symptoms [22–29], and its effectiveness varies depending on the time of intervention, with little effect on ventilated COVID-19 patients [30]. Molnupiravir and favipiravir offer oral convenience but have shown limited clinical effectiveness [31–34]. Paxlovid is an effective oral therapy, however, the combination drug ritonavir, strongly inhibits CYP3A metabolism, creating substantial drug–drug interaction risks with commonly prescribed medications [35]. While the recently approved drug, ensitrelvir, an orally active non-covalent main protease (M^pro^) inhibitor, does not require ritonavir co-administration, its cost (circa US $1400 for a five-day regimen) is unaffordable to many parts of the world (c.f. India’s GDP per capita of circa US $2800). Besides, it exerts a strong inhibitory effect on cytochrome P450 and other key drug transporters, such as P-glycoprotein [36], which may alter the metabolism of other drugs [37]. Co-administration with susceptible substrates/inducers may alter their metabolism and/or transport, thereby increasing systemic drug exposure and the risk of concentration-dependent adverse effects. This may restrict the concomitant use of ensitrelvir (especially drugs with narrow therapeutic index) – an important consideration in patients receiving treatments for multiple diseases. Therefore, there remains a need for the development of safe, accessible, affordable and mechanistically distinct broad-spectrum agents capable of influencing viral and host-mediated processes associated with COVID-19, PASC (post-acute sequalae of COVID-19, also known as long COVID), and any future “Disease X”.

Drug repurposing is a practical strategy to address some of these limitations by leveraging clinically approved compounds with known therapeutic, toxicological and safety profiles, thereby reducing developmental costs and accelerating early drug discovery [38]. To systematically identify such candidates with potential anti-SARS-CoV-2 activity, we previously established a systematic pipeline that included the development of a curated database (CoviRx) [39], followed by strategic down-selection of molecules based on pharmacokinetic and pharmacodynamic profiles to 214 drugs [40] and subsequent final selection and *in vitro* testing of the top 12 hit molecules in organoid models [41]. In our *in vitro* analysis, we identified three drugs with anti-SARS-CoV-2 activity, *viz.* fluvoxamine (selective serotonin reuptake inhibitor or SSRI), everolimus (mechanistic target of rapamycin inhibitor or mTOR), and rolapitant (neurokinin-1 receptor inhibitor or NK-1), however none of these demonstrated sufficient antiviral activity for use as an antiviral monotherapy [41]. We extended our search to identify which other drugs among the 214 warranted further *in vitro* evaluation, carried out an *in silico* screening of the down-selected molecules across various SARS-CoV-2 targets, and identified four candidates (alendronate, cromolyn, natamycin and treprostinil) for *in vitro* testing [38]. Alendronate is a bisphosphonate (BP) widely used in the treatment of osteoporosis [42,43], while cromolyn is a mast cell stabilizer used in the treatment of allergic rhinitis and asthma [44]. Natamycin is a naturally occurring anti-fungal agent and treprostinil is a synthetic prostacyclin analog used for treating pulmonary arterial hypertension [45,46].

Among these, alendronate gained particular attention in our study because the existing literature reported its binding to the conserved Nidovirus RdRp associated nucleotidyltransferase (NiRAN) domain of the coronavirus, providing an important putative mechanistic foundation for further investigation [47]. This mechanistic evidence motivated a focused virtual screening of all nitrogen containing bisphosphonates (N-BPs) in the ChEMBL database against the SARS-CoV-2 RdRp protein. We identified seven promising BPs, two of them closely resembling minodronic acid (CHEMBL608526) and zoledronic acid (CHEMBL98211) [48] (**Figure 1**). In parallel, a study led by the Harvard Medical School on 450,366 COVID-19 patients who did not use BPs vs 450,366 COVID-19 patients with prior use of alendronate/alendronic acid or zoledronate/zoledronic acid found that the latter BP users had lower odds ratios (OR) of COVID-19 infection (OR = 0.22; 95% CI:0.21–0.23), diagnosis (OR = 0.23; 95% CI:0.22–0.24) and related hospitalization (OR = 0.26; 95% CI:0.24–0.29) [43]. Our previous papers [38,48] nicely complemented this Harvard study by providing the missing mechanistic explanation for Harvard’s compelling real-world evidence [43] which is also supported by further evidence [49,50]. While other studies did not observe any significant benefit against COVID-19 [42,51,52], we believed that the overarching evidence was still in favor of BPs and that it was worthwhile investigating if they had an antiviral effect. Observational studies cannot establish whether any protective association results from a direct antiviral effect of BPs or from indirect effects related to host physiology, comorbidities, concomitant therapies, or other factors. Therefore, considering these uncertainties, and our recent findings on fluvoxamine, which demonstrate both antiviral and host-associated mechanisms, there is a compelling need to evaluate the antiviral activity of BPs and elucidate their underlying mechanisms of action.

**Figure 1:**
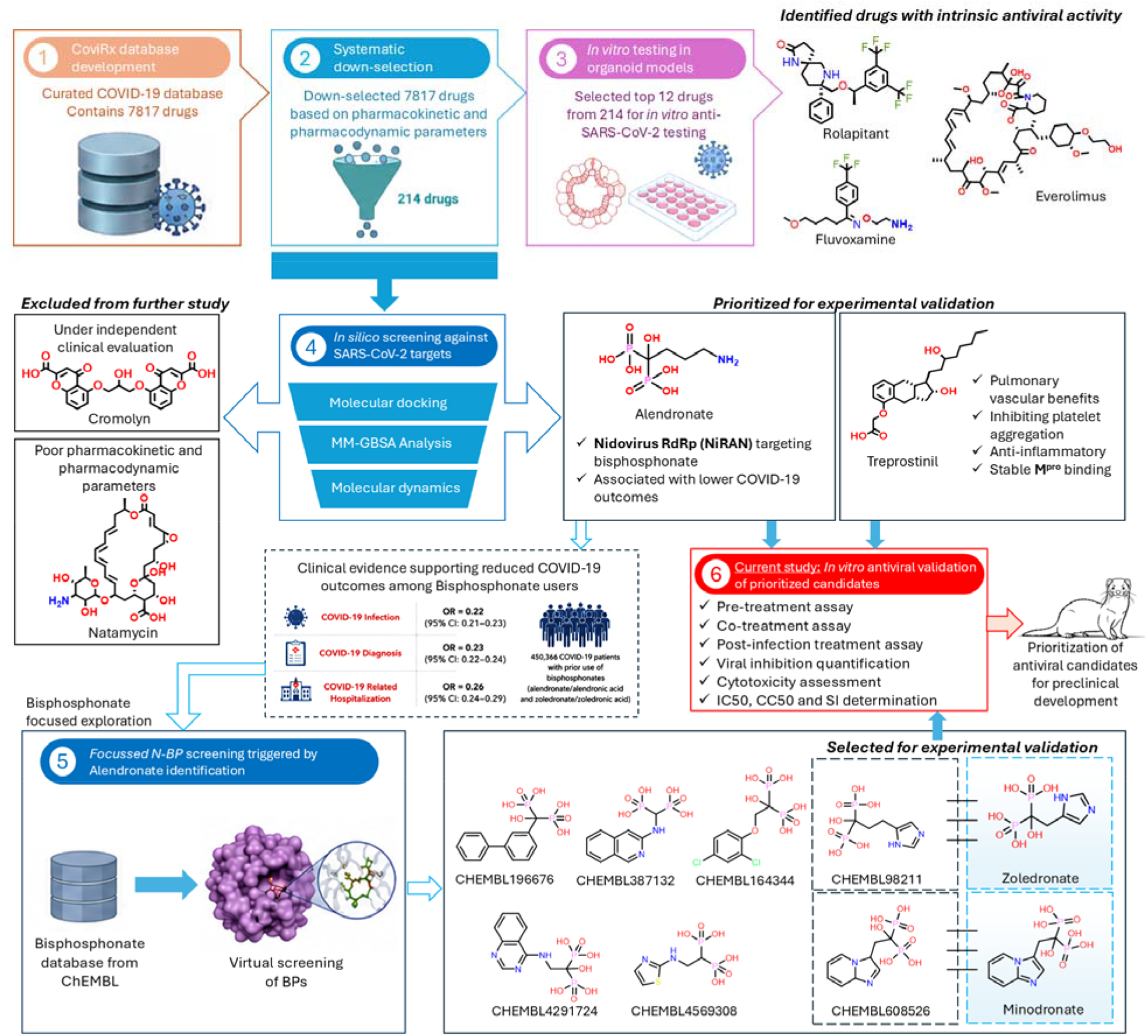
Integrated workflow for identification and experimental validation of repurposed SARS-CoV-2 inhibitors through CoviRx screening and bisphosphonate-focused drug discovery

## Materials and Methods

The Omicron variant of SARS-CoV-2 isolate (PQ.8.1) was isolated at the Australian Centre for Disease Preparedness (ACDP; Geelong, VIC, Australia) from a nasopharyngeal swab obtained by Barwon Health (the local health authority). Swab samples in virus transport medium were diluted 1:2 in PBS containing 4x antibiotic-antimycotic (ThermoFisher Scientific; Scoresby, VIC, Australia) and filtered through a 0.45µm filter to remove cellular contamination. 1ml of sterilized inoculum was added to rinsed Vero E6 cells (American Type Culture Collection; Manassas, VA, USA) in a T-75 flask and incubated for 1hr at 37°C/5% CO_2_. After the incubation, the cells were rinsed with PBS and 10ml maintenance medium (Dulbecco’s Minimum Essential Medium (DMEM) containing 1% FBS, 2mM GlutaMAX supplement, 100U/mL penicillin, and 100 μg/mL streptomycin (all components from ThermoFisher Scientific)) was added. Flasks were returned to the incubator and checked daily for the development of cytopathic effect (CPE). Once CPE had developed, supernatant samples were collected centrifuged at 2,000 x g for 10 min to clarify, harvested and stored in 1 mL aliquots at −80 °C.

Working stocks were grown in T-150 flasks containing Vero E6 cells, with isolated virus material diluted to 1:100 in Dulbecco’s Minimum Essential Medium (DMEM) added with 2% FBS, 2mM GlutaMAX supplement, 100U/mL penicillin, and 100 μg/mL streptomycin (all components from ThermoFisher Scientific). Diluted inoculum (5 mL) was used to inoculate Vero E6 cells for 1 hr at 37°C/5% CO_2_ before 35ml additional media was added to the flask. The flasks were incubated for 48 h before supernatant was centrifuged at 2,000 x g for 10 min to clarify, harvested and stored in 1 mL aliquots at −80 °C.

Identity of virus stocks was confirmed by next-generation sequencing using a MiniSeq platform (Illumina, Inc.; San Diego, CA, USA). In brief, 100 µL cell culture supernatant from infected Vero E6 cells (virus isolation and stock culture) was combined with 300 µL TRIzol reagent (ThermoFisher Scientific) and RNA was purified using a Direct-zol RNA Miniprep kit (Zymo Research; Irvine, CA, USA). Purified RNA was further concentrated using an RNA Clean-and-Concentrator kit (Zymo Research), followed by quantification on a DeNovix DS-11 FX Fluorometer. RNA was converted to double-stranded cDNA, ligated then isothermally amplified using a QIAseq FX single cell RNA library kit (Qiagen; Hilden, Germany). Fragmentation and dual-index library preparation was conducted with an Illumina DNA Prep, Tagmentation Library Preparation kit. Average library size was determined using a Bioanalyser (Agilent Technologies; San Diego, CA, USA) and quantified with a Qubit 3.0 Fluorometer (Invitrogen; Carlsbad, CA, USA). Denatured libraries were sequenced on an Illumina MiniSeq using a 300-cycle Mid-Output Reagent kit as per the manufacturer’s protocol. Paired-end Fastq reads were trimmed for quality and mapped to the published sequence for the SARS-CoV-2 reference isolate Wuhan-Hu-1 (Ref Seq: NC_045512.2) using CLC Genomics Workbench version 21 from which a consensus sequence was generated. Stocks were confirmed to be free from contamination by adventitious agents by analysis of reads that did not map to SARS-CoV.

### Test and control drug molecules

In accordance with the aforesaid findings and Figure 1, we have selected three commercially available BP molecules — alendronate/alendronic acid from the initial *in silico* screening [38]; minodronate/minodronic acid and zoledronate/zoledronic acid from the subsequent *in silico* BPs screening [48]— to evaluate their antiviral efficacy *in vitro*. We have retained treprostinil from our initial screening [38], given its beneficial effects for COVID-19 patients through improving pulmonary vascular oxygenation, inhibiting platelet aggregation and anti-inflammatory property [45,46], and our recent findings showing stable interaction profile with SARS-CoV-2 M^pro^ [38]. The other two drugs, cromolyn and natamycin were excluded as the former drug had been under independent clinical investigation [53], while the latter drug has poor pharmacokinetic properties that may restrict systemic therapeutic effect. Thus, the present study experimentally evaluated the antiviral activity of key BPs identified through our repurposing pipeline to determine if the reported clinical association of BPs with COVID-19 outcomes could be supported by any intrinsic antiviral activity, or if these effects are due to other mechanisms of action. For control drugs, we retained remdesivir and nirmatrelvir used in our previous paper [41], and substituted molnupiravir for ensitrelvir, as these drugs are the current standard of care.

### Drug stock solutions

Alendronate, ensitrelvir, nirmatrelvir, remdesivir and treprostinil were procured from Med Chem Express (Monmouth Junction, NJ, USA), while zoledronate monohydrate and minodronate were procured from BLD Pharmatech Ltd. All compounds were obtained in powder form and reconstituted using an appropriate solvent prior to use. Alendronate, minodronate and zoledronate were dissolved in 1 M NaOH due to limited solubility in aqueous buffers and DMSO. Ensitrelvir, nirmatrelvir and treprostinil and were dissolved in DMSO. All drugs were prepared as 25 mM stock solutions. Remdesivir was obtained as a pre-prepared 10 mM stock solution (SelleckChem; Houston, TX, USA). Drug working stocks were prepared inside a biosafety cabinet and sterilized by filteration through 0.22 µm syringe filters before aliquoting into single-use microcentrifuge tubes. All aliquots were stored at −80°C until use. Equivalent solvent-only control groups containing DMSO or NaOH were included in all experiments.

### Cytotoxicity assay

The cytotoxicity of the selected compounds was evaluated prior to antiviral testing using the Thermo Scientific CyQUANT LDH Cytotoxicity Assay kit. VeroE6 cells were seeded into 96-well tissue culture plates and incubated overnight at 37°C with 5% CO to allow cell attachment. These cells were treated in triplicate with serial dilutions of the selected drugs and the standards at concentrations of 25, 10, 4, 1, 0.4, and 0.08 µM. Four main control conditions — blank, spontaneous LDH, maximum LDH, positive LDH and solvent only, were included. Plates were incubated for 6 and 48hr at 37°C/5% CO2 to allow determination of acute and non-acute drug-induced cytotoxicity, respectively. Post incubation, 10µl of lysis buffer was added to the maximum LDH release wells, followed by a 45 min incubation. Next, 50 µL of culture supernatant from each well was transferred into black 96-well fluorescence plates followed by the addition of reagent stock solution as per the manufacturer’s protocol, and fluorescence was measured using a microplate reader at excitation and emission wavelengths of 560 and 590 nm, respectively. Cytotoxicity was calculated as a percentage of the maximum LDH release observed in the controls using the following formula.

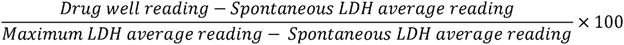

### SARS-CoV-2 antiviral assay

The antiviral activity of the selected compounds was evaluated against a clinical SARS-CoV-2 isolate (PQ.8.1) in VeroE6 cells in a 96-well plate format. Test and control drugs were diluted in DMEM supplemented with 10% FBS, 2 mM Glutamine supplement and 10mM HEPES, to final concentrations of 25, 10, 4, 1, 0.4, and 0.08 µM. Drug treatment was performed using a three-stage protocol consisting of pre-treatment, co-treatment during infection, and post-infection treatment. In the pre-treatment stage, six replicate dilution series per sample were dispensed into wells of a 96-well plate (50 µL per well) containing pre-plated 4 × 10^4^ Vero E6 cells/well. Plates were incubated at 37°C/5% CO_2_ for 1h at 37°C and 5% CO_2_. After pretreatment, the medium is replaced with virus inoculum (5x10^4^ TCID_50_/ml, or 5x10^3^ TCID_50_/100µl final concentration) along with the corresponding drug concentrations and incubated for 1h under same conditions as pretreatment to allow the virus to adsorb. After viral adsorption (post-infection), the inoculum was replaced with 200 µL of the medium containing the corresponding drug concentration and incubated for 48 h at 37°C in a humidified 5% CO atmosphere. After incubation, the culture supernatants from each well were collected into sterile pre-labelled Sarstedt tubes and stored at −80°C until downstream viral titration analysis (**Figure 2**). Mock-infected controls, virus-only controls, and solvent controls were also included in each experiment. Antiviral activity was determined by comparing viral titters from treated wells with untreated virus-infected control wells. Reduction in viral replication relative to the virus control was considered indicative of antiviral activity.

**Figure 2:**
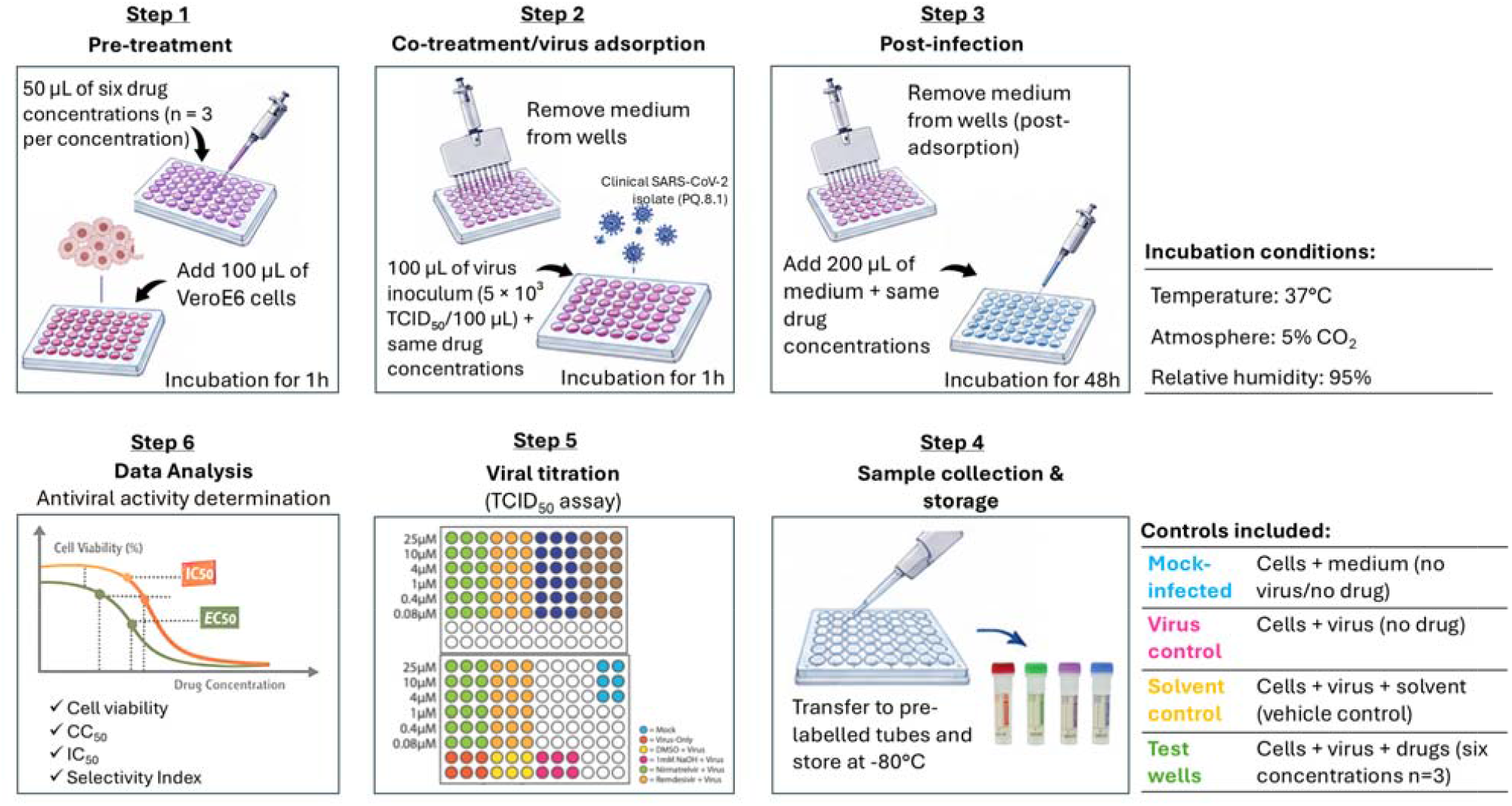
Schematic representation of the experimental workflow for *in vitro* evaluation of antiviral activity of test compounds against SARS-CoV-2 in Vero E6 cells

### TCID_50_ assay

Virus titres were determined by endpoint dilution assay using Vero E6 cells grown in DMEM, 10% FBS, Glutamine and HEPES. Cells were seeded into clear 96-well plates to reach confluency 24 hours prior to infection. Each sample was assessed across serial ten-fold dilutions using 4 replicate wells per sample. Following preparation of the serial dilutions, culture medium was removed from the assay wells and replaced with the corresponding virus dilution. Plates were incubated under standard cell culture conditions for 72 hours. Cells were fixed with 4% PFA and analysed for CPE. Infectious virus titres were calculated across the serial dilution series using the TCID50 Spearman-Kärber method.

### Immunofluorescent staining

Cells were permeabilised with 0.1% TritonX for 10 minutes at RT, and subsequently blocked using 0.5% bovine serum albumin (BSA) in phosphate-buffered saline (PBS) for 30 minutes at RT. SARS-CoV-2 nucleocapsid protein was detected using a mouse monoclonal anti-SARS/SARS-CoV-2 nucleocapsid antibody (clone 05; MA5-29981; Thermo Fisher Scientific), incubated overnight at 4C, followed by 3 washes with PBS and a 1 hour incubation at room temperature in Alexa Fluor 488-conjugated goat anti-mouse IgG secondary antibody (A-11001; Thermo Fisher Scientific) followed by a 10 minute incubation in 5mg/ml DAPI (Cat#62248; Thermo Fisher Scientific) at RT. Images were taken at 20x Magnification on a ZEISS LSM800 confocal microscope.

## Results and discussion

### Cytotoxicity of selected compounds

The cytotoxicity of the selected compounds was evaluated in Vero E6 cells prior to assessment of antiviral activity. The cytotoxicity assay was performed at concentrations ranging from 0.08 to 25 µM, as described in the materials and methods section, with 6 and 48hr cellular incubations to assess potential acute and non-acute toxicity. All test and control compounds showed minimal cytotoxicity at concentrations up to 25 µM in the LDH release assay. These findings indicated that the concentrations selected for subsequent antiviral evaluation were well tolerated by Vero E6 cells and that any reduction in viral replication could be assessed without an apparent confounding effect of drug-induced cellular toxicity.

### Evaluation of antiviral activity by immunofluorescence

The antiviral activity of all test and control drugs against SARS-CoV-2 PQ.8.1 was initially evaluated by immunofluorescence microscopy. Mock-infected cells showed predominantly blue nuclear staining with minimal green fluorescence, whereas cells infected in the absence of treatment showed prominent green fluorescence, consistent with extensive viral infection. Treatment with the test compounds did not result in an apparent reduction in the green fluorescence signal compared with the virus-only control across the concentrations examined (Figure 3). In contrast, treatment with reference antiviral compounds (ensitrelvir, nirmatrelvir, remdesivir) resulted in a marked reduction in the green fluorescence signal, with staining patterns approaching those observed in mock-infected cells (supplementary figure 1). These observations indicate that the experimental system was responsive to established antiviral agents, whereas the investigational compounds did not produce an appreciable reduction in the virus-associated fluorescence signal under the conditions tested.

**Figure 3:**
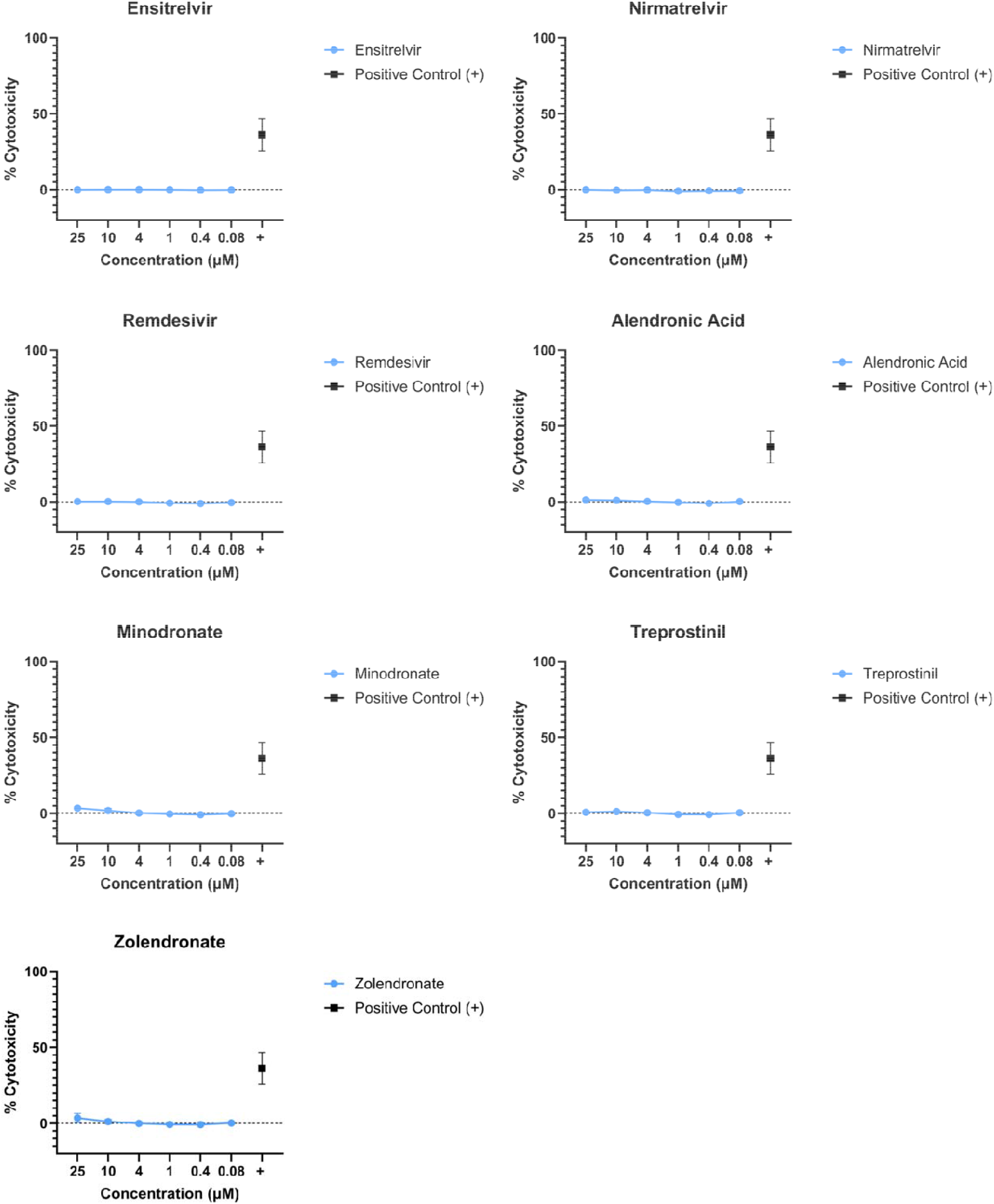
Cytotoxicity of selected compounds in Vero E6 cells. Cytotoxicity of Ensitrelvir, Nirmatrelvir, Remdesivir, Alendronic acid, Minodronate, Treprostinil, and Zoledronate was assessed by LDH release across 0.08–25 µM at either 6 or 48 hours. Data are mean ± SD. All compounds showed minimal or no detectable cytotoxicity, supporting their use in subsequent antiviral assays

### Effect on infectious viral titres

The effect of alendronate, minodronate, and zoledronate on infectious virus titres was determined by TCID assay. Viral titres remained broadly comparable across the concentration range of 0.08–25 µM for all three compounds (Figure 4). None of the BPs produced a consistent concentration-dependent reduction in infectious viral titre, with titres remaining approximately around 5 log TCID /mL across the concentration range examined. A similar pattern was observed following treatment with treprostinil for which viral titres remained approximately within the 5–6 log TCID /mL range. The small variations in viral titre observed between individual concentrations did not follow a consistent dose-response pattern and therefore did not provide evidence of concentration-dependent inhibition of viral replication. Collectively, the TCID data support the immunofluorescence findings and indicate that BPs and treprostinil do not demonstrate significant antiviral activity against SARS-CoV-2 PQ.8.1 under the experimental conditions employed. While these negative results are disappointing for treprostinil, they are not unexpected for the three BPs because there has been no studies or suggestions to date indicating direct antiviral property and the plasma concentrations for these compounds are quite low — 161 nM for alendronate [54], 4.36 nM for minodronate [55] and 1,134 nM for zoledronate [56]. The fact that the present study only used a single SARS-CoV-2 isolate is theoretically a limitation, however, we are confident that PQ.8.1 is a representative isolate given its isolation from a community SARS-CoV-2 infection and our long experience in the fight against COVID-19 since early 2020 [5]. Factors such as cellular uptake, intracellular concentration, compound stability, target accessibility and the contribution of host-cell pathways could also have influenced the observed antiviral phenotype.

**Figure 4.**
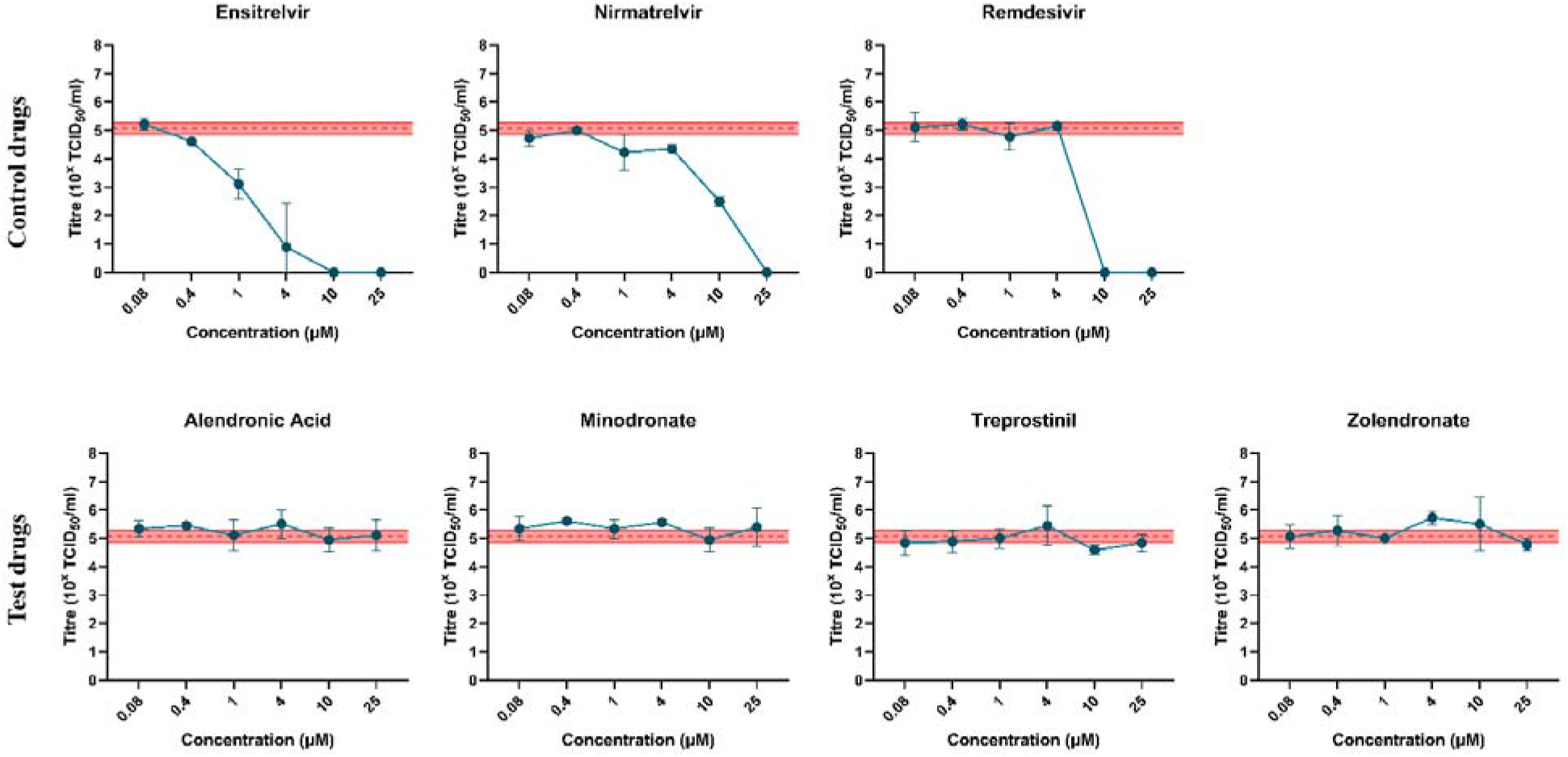
Effect of bisphosphonates on infectious SARS-CoV-2 titre. Infectious viral titres in culture supernatants following treatment with alendronate, ensitrelvir, minodronate, nirmatrelvir, remdesivir, treprostinil and zoledronate.at concentrations ranging from 0.08 to 25 µM. Viral titres were determined by TCID assay following 48 h of infection.

### Other potential mechanisms of action

With direct antiviral activity ruled out by this study, any beneficial effects such as the one described by the Harvard study [43], are likely to involve host-directed or immunomodulatory, anti-inflammatory or other mechanisms of action. Our previous computational studies suggested a potential virus-mediated mechanism of action, i.e the three BPs evaluated in this paper (alendronate, minodronate, zoledronate) are potential binders of the conserved NiRAN domain-associated region of the coronavirus polymerase, providing a potential mechanistic basis for subsequent evaluation of related N-containing BPs [38,48]. Future studies should investigate host-mediated mechanisms, for instance it has been suggested that the effects on γδ-T-cell populations and protein prenylation pathways could potentially influence host responses to viral infection [57]. Moreover, modulation of cathepsin-associated pathways could represent another potential host-directed mechanism.

Lysosomal cysteine protease, particularly cathepsin B and L, plays an important role in the entry of SARS-CoV-2 and facilitates the fusion and release of viral genome into the host cell [58][59]. An increase in the cathepsin L (CTSL) expression showed enhanced SARS-CoV-2 infection, whereas its knockdown or pharmacological inhibition impaired spike processing, viral entry, and infection [60]. Additionally, cathepsins B and L, and other cysteine cathepsins, including K, S, and V, have been reported to cleave the S protein at multiple sites suggesting their vital role in viral entry and pathogenesis [59]. Interestingly, cathepsin activity is also closely linked to inflammatory responses, cell death, extracellular-matrix remodeling and tissue injury, which are of interest because N-containing BPs have been reported to suppress cathepsin K (CTSK) expression in osteoclasts [61–63]. Although CTSK is associated with osteoclastic bone resorption, its reported ability to process the SARS-CoV-2 spike protein, together with its involvement in inflammatory and tissue-remodeling processes, raises the possibility that BP-mediated modulation of cathepsin-associated pathways could influence host responses during infection. The ability of pharmacological cathepsin inhibitors to reduce SARS-CoV-2 entry and infection in experimental models does support cathepsins as potential host-directed therapeutic targets [64]. However, modulation of CTSK or other cathepsins may not always be beneficial because cathepsins also perform several physiological functions, and any severe dysregulation could contribute to pathological inflammation and tissue remodeling responses. Taken together, these observations suggest that modulation of cathepsin-dependent pathways could contribute to some of the observed effects associated with BPs in COVID-19. However, a direct relationship between BP treatment, cathepsin modulation, and altered SARS-CoV-2 infection has not yet been established and requires further exploration including experimental validation.

### Conclusions

Molecular docking and related *in silico* approaches can identify compounds with potential interactions with viral targets, but such interactions do not necessarily translate into intracellular target engagement or inhibition of viral replication, thus demonstrating the importance of experimental validation as outlined in this study. Although there was no detectable cytotoxicity up to 25µM in VeroE6 cells, the investigational compounds (alendronate, minodronate, zoledronate, as well as treprostinil) did not demonstrate an appreciable antiviral effect against SARS-CoV-2 PQ.8.1 in comparison to control drugs (ensitrelvir, nirmatrelvir and remdesivir). These negative results are disappointing for treprostinil, but they are in line with expectations for BPs. While this study does not preclude efficacy against other viruses in other physiologically relevant cell systems, it is reasonable to conclude that the test compounds are unlikely to be direct antiviral agents for betacoronaviruses. Even though they lack antiviral activity, if BPs protect through other potential mechanisms of action such as cathepsins or RdRp, then such effects could inflate the efficacy of vaccines and drugs tested on those users. Therefore, studies on COVID-19, PASC and future “Disease-X” should be appropriately designed, given that some of these could involve conserved regions across RNA viruses.

## Abbreviations

BPs: bisphosphonates
COVID-19: coronavirus disease 19
CTSK: cathepsin K
CTSL: cathepsin L
CYP3A: cytochrome member 3A
DMSO: dimethyl sulfoxide
DMEM: Dulbecco’s Modified Eagle’s Medium
HIV: human immunodeficiency virus
HEPES: (4-(2-hydroxyethyl)-1- piperazineethanesulfonic acid) buffer
LDH: Lactate Dehydrogenase
Mpro: main protease
mTOR: mechanistic target of rapamycin
N-BPs: nitrogen containing bisphosphonates
NK-1: Neurokinin-1
NiRAN: Nidovirus RNA binding domain
PASC: post-acute sequalae of COVID-19
PQ.8.1: SARS-CoV-2 Omicron-derived sublineage descending from the JN.1 and NB.1.8.1 lineage
RNA: ribonuclease protein
RdRp: RNA dependent RNA polymerase
SARS: severe acute respiratory syndrome
SARS-CoV-2: SARS coronavirus 2
TCID50: Tissue Culture Infectious Dose 50
Vero E6: kidney epithelial cells derived from an African green monkey
Wuhan-Hu-1: original reference genome strain of SARS-CoV-2

## Acknowledgements

The authors thank Professor Trevor W. Drew OBE for his comments. MMUR and SO are grateful for resources received in kind from BITS Pilani and CSIRO. AA is partially supported by CSIR-National Laboratories Scheme under the ULIP sub-scheme MLP002605. SSV acknowledges funding from BITS Pilani 1991 Alumni and the Royal College of Physicians and Surgeons of Glassgow for the Triannual Travel Medicine Research Fellowship Award (2024-27) to pursue Disease-X research. He is grateful to the United Sates Food and Drug Administration (US FDA) Medical Countermeasures Initiative for funding previous work (Contract Number: 75F40121C00144) which directly led to this study, and for inviting him to present the preliminary findings at Silver Spring (MD) on May 31, 2024.

## Author contributions

**Conceptualization**: SSV; **Methodology**: AJM, MMUR, SSV; **Software**: MMUR; **Validation**: AJM, MMUR; **Formal analysis**: AA, AJM, MMUR, SO, SSV, TWD; **Investigation**: MMUR, SO; **Visualization**: AJM, MMUR; **Resources**: AJM, SM, SSV; **Funding acquisition**: AJM, SM, SSV; **Data curation**: AJM, MMUR, SO, SSV; **Writing original draft**: AA, AJM, MMUR, SSV, TWD; **Writing-review and editing**: AA, AJM, KVGCS, MMUR, SM, SO, SSV, TWD; **Project administration**: AJM, SSV; **Supervision**: AJM, KVGCS, SM, SSV.

## Conflicts of Interest

SSV is Moderna Respiratory Portfolio Advisory Board Member. All other authors declare no conflict of interest. This article reflects the views of the authors and does not represent the views or policies of affiliating institutions or funding agencies.

## Data Availability Statement

Any underlying data not presented can be provided by the corresponding author upon reasonable request.

## Funding

Not applicableInformed Consent Statement

Not applicable

## Institutional Review Board Statement

Virus isolations from human diagnostic samples were performed with approval from the CSIRO Health and Medical Human Research Ethics Committee (CHMHREC; Approval Number: 2025_005_LR).

