## Supplementary file 1 for "Experimental validation of computationally prioritized bisphosphonates reveals no direct *in vitro* antiviral activity against SARS-CoV-2"

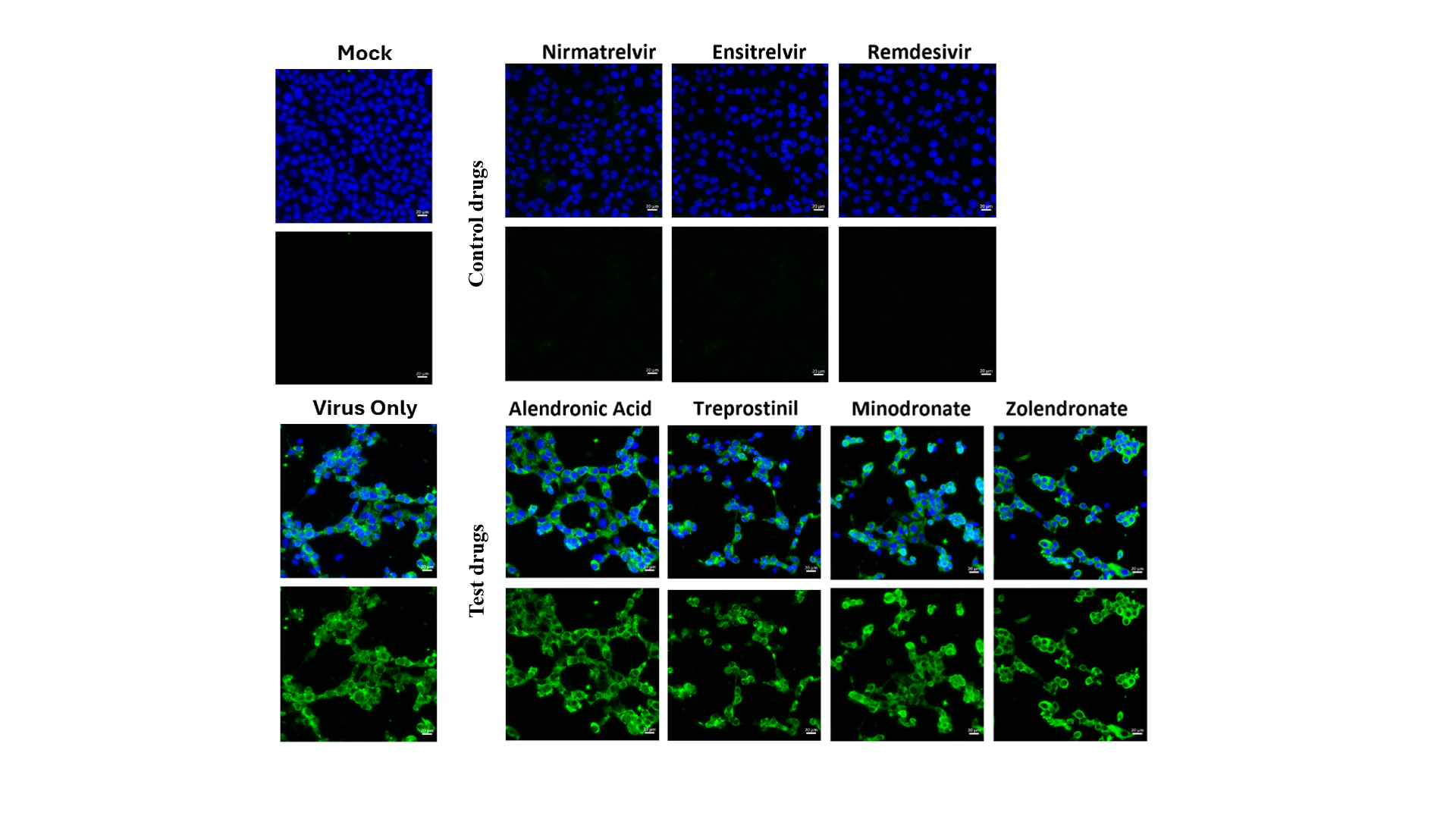

**Supplementary figure 1. Immunofluorescence analysis of SARS-CoV-2 infection following treatment with selected compounds.** Representative immunofluorescence images of Vero E6 cells infected with SARS-CoV-2 PQ.8.1 following treatment with alendronate, ensitrelvir, minodronate, nirmatrelvir, remdesivir, treprostinil and zoledronate. Mock-infected and virus-only conditions were included as negative and infection controls, respectively. Blue fluorescence represents nuclear staining, while green fluorescence represents the virus-associated immunofluorescence signal.
